# Systematic Discovery of Phage Genes that Inactivate Bacterial Immune Systems

**DOI:** 10.1101/2024.04.14.589459

**Authors:** Shinjiro Ojima, Aa Haeruman Azam, Kohei Kondo, Wenhan Nie, Sai Wang, Kotaro Chihara, Azumi Tamura, Wakana Yamashita, Tomohiro Nakamura, Yo Sugawara, Motoyuki Sugai, Bo Zhu, Yoshimasa Takahashi, Koichi Watashi, Kotaro Kiga

## Abstract

Bacteria have developed numerous defense systems to counter phage infections. However, the extent to which phages possess countermeasures against these defense systems remains unclear. In this study, we combined a phage gene knockout library with a defense system library to analyze the mechanisms by which phages counteract bacterial defense systems. After attempting gene deletions of 105 open reading frames (ORFs) in the DruSM1 phage (ΦDruSM1), we successfully generated 73 different ORF knockout phages. By infecting this library with bacteria harboring defense system expression plasmids, we identified inactivators of Druantia type I (Druad1), Brex type I, AVAST type III, Sir2+HerA, DUF4297+HerA, and hhe, as well as an activator of Retron Ec86, in a single phage genome. Synthetic phages incorporating Druad1 effectively eradicated *Escherichia coli* harboring the robust Druantia type I defense system by altering DNA methylation at m6A sites of the phage. This study highlighted the prevalence of various antidefense mechanisms employed by phages to overcome bacterial defense systems.

## Introduction

Bacteria have developed diverse antiphage immune systems to survive phage infections. Among these, systems such as restriction modification and CRISPR-Cas have been recognized as mechanisms for cleaving DNA of targeted phages^1^; however, recent research has revealed that bacteria utilize a broader range of defense mechanisms, and more than 100 different defense mechanisms have been reported^2,3^. These bacterial defense systems have forced bacteria-infecting phages to evolve various evasion strategies, including anti-Retron^4^, anti-Thoeris, anti-Gabija^5,6^, anti-AVAST^7^, anti-CBASS, anti-Pycsar^8,9^, and anti-BREX^10-12^. Phages also use tRNA as an antidefense against retron Ec78^4^. For further understanding of defense systems and enhancing the effectiveness of phage therapy hindered by defense systems, elucidating antidefense mechanisms is crucial. However, the functionality of antidefense mechanisms remains largely unknown, representing an important unsolved issue in the field.

The limited discovery of defense systems can be attributed to the lack of established strategies or the requirement for technically challenging experiments. The anti-CBASS gene *acb1* and anti-Pycsar gene *apyc1* were discovered by comparing the cyclic nucleotide-degrading activity following phage infection^8^. The anti-Thoeris gene *tad1* was discovered by comparing the genome sequences of similar phages^5^. In both studies, candidate anti-defense genes were cloned into plasmid vectors and coexpressed with their respective defense systems in host bacteria to determine their neutralizing activity against the bacterial defense system during phage infection. However, phage-derived genes are often toxic to host bacteria, and stable expression of anti-defense genes in plasmids is challenging^13^.

We previously identified antidefense genes by studying naturally occurring phage gene deletion strains^4^. In this study, we attempted to identify antidefense genes by artificially generating phage gene deletion mutants, in which each open reading frame (ORF) of the phage is deleted one by one^14,15^.

## Results

### Construction of a phage gene knockout library

The generation of gene knockout libraries is extremely useful for studying the function of genes in organisms^16^. However, a systematic phage knockout library has not yet been constructed. First, we focused on ΦDruSM1. This phage belongs to the Quenovirinae phage family, with a genome size of 60 kb, which is reasonable for constructing a knockout library using in vitro synthesis methods (Figure 1A)^17,18^. The phage genome was amplified by PCR using primers specifically designed to delete a particular gene, and the resulting fragments were assembled to generate a circular genome. This artificially synthesized genome was electroporated into *E. coli* HST08, and the phage was rebooted. The number of plaques generated by rebooting the synthesized phage varied greatly depending on the deleted gene (Figure 1A, B). If plaque formation of the gene-deleted phage was less than 10 after rebooting, the deleted gene was considered essential for the phage. The reason for setting the threshold at 10 was that even when the capsid genes, which are already known as essential genes, were deleted, several to around 10 plaques were still formed. Consequently, in ΦDruSM1, 32 genes were assumed to be essential, whereas 72 genes were assumed to be nonessential genes. The genes determined to be primarily essential are terminase, capsid, tail structure, and nucleotide metabolism (Figures 1C and S1B). This is consistent with a previous study reporting that structural genes and genes involved in DNA replication in phage are essential^19^. Overall, we successfully generated ORF knockout mutants of 72 nonessential genes and used them in further experiments.

**Figure 1.**
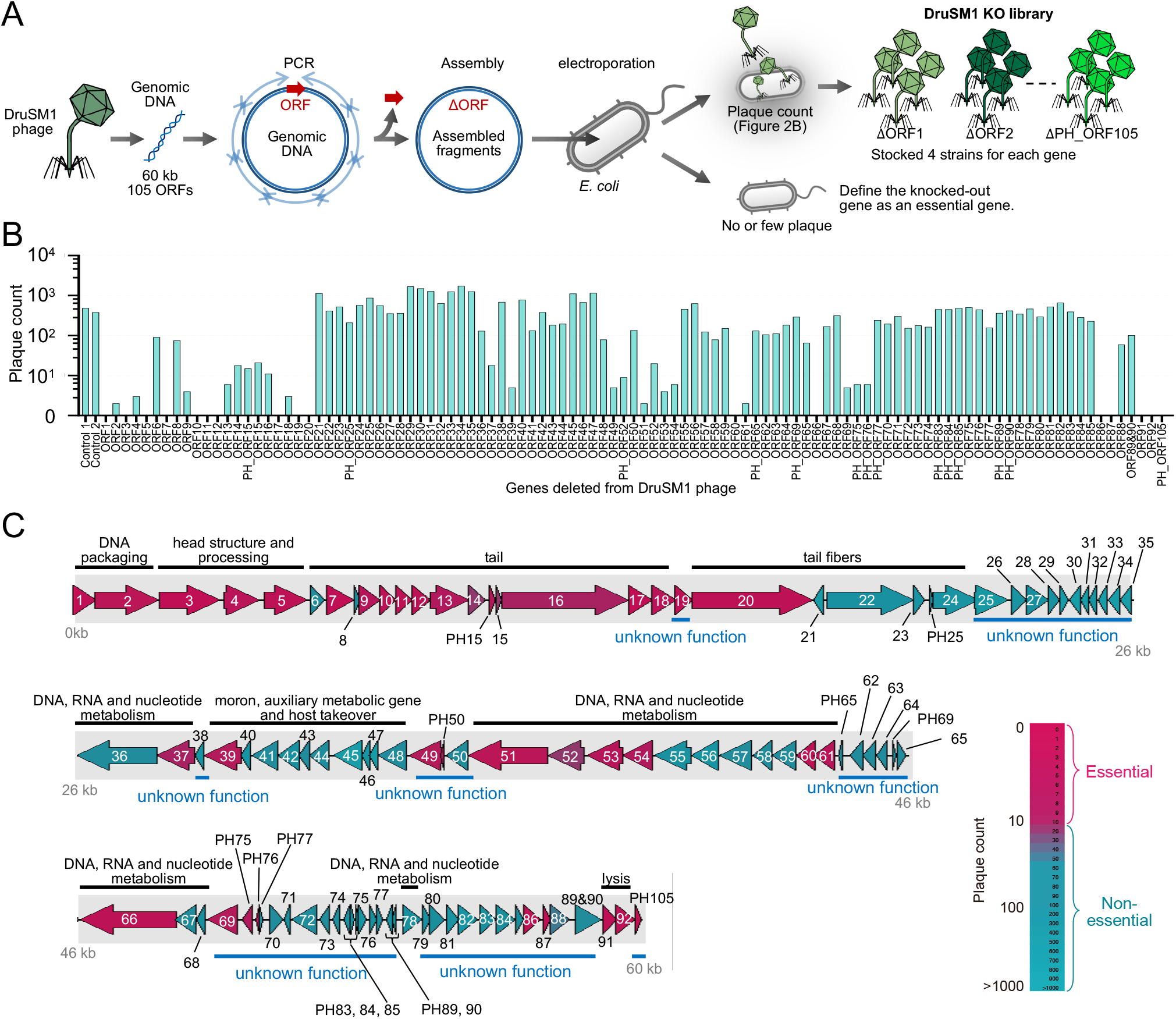
Construction of gene deletion library of ΦDruSM1 phage (A) Schematic diagram of the construction of the phage gene deletion library. Deletion of ORFs was performed by excluding PCR fragments from the genome of ΦDruSM1 phage, assembling the fragments, and rebooting the phage using *E. coli*. (B) The number of plaques appearing on the plate after rebooting each ORF deletion phage were counted. (C) Genome map of ΦDruSM1 phage ORFs colored by the number of plaques obtained in (B). ORFs that could not be deleted are colored wine red, whereas those that could be deleted are colored turquoise blue. ORF annotations were done using pharokka.

### Identification of phage-derived genes that alter the activity of bacterial defense systems

To investigate the phage genes affecting the sensitivity to bacterial defense systems, the constructed gene-deletion phage library was used to infect *E. coli* DH10B cells harboring the defense system library (Figure 2A)^20^. As synthetic phages can unintentionally acquire genetic mutations during construction, four independent synthetic phage strains were constructed for each ORF-deleted phage. Accordingly, 19 types of gene-deleted phages for which the infection efficiency was reduced by more than 100-fold in at least three independent experiments were obtained (Table 1). Seven types of gene-deleted phages were identified with a 10- to 100-fold decrease. Four types of gene-deleted phages exhibited a 100-fold or greater increase in efficiency. Genomic loss in the mutant phages with altered efficiency of plaque formation (EOP) was confirmed by PCR (Figure S1A).

**Table 1.**
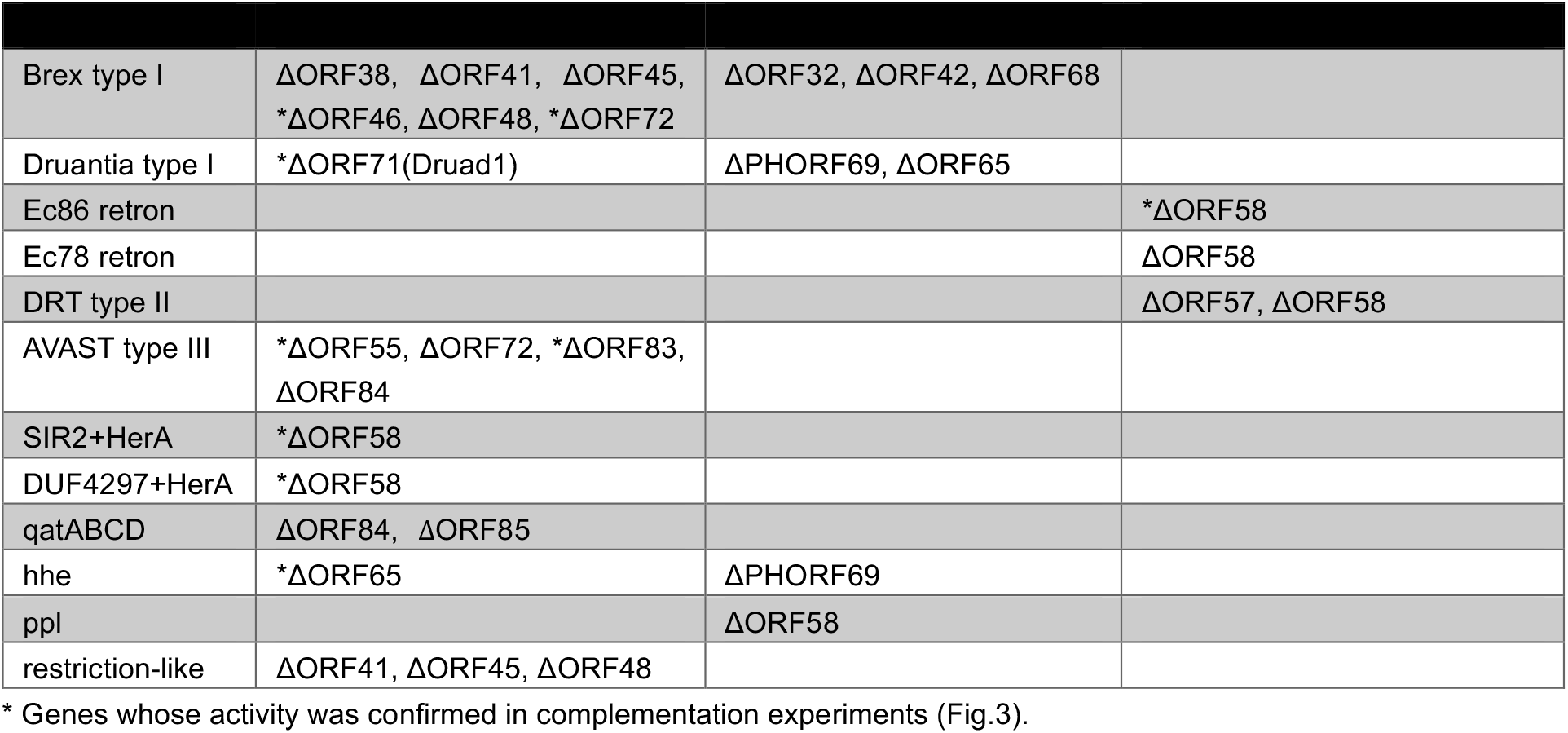
List of deleted genes with altered susceptibility to defense systems.

**Figure 2.**
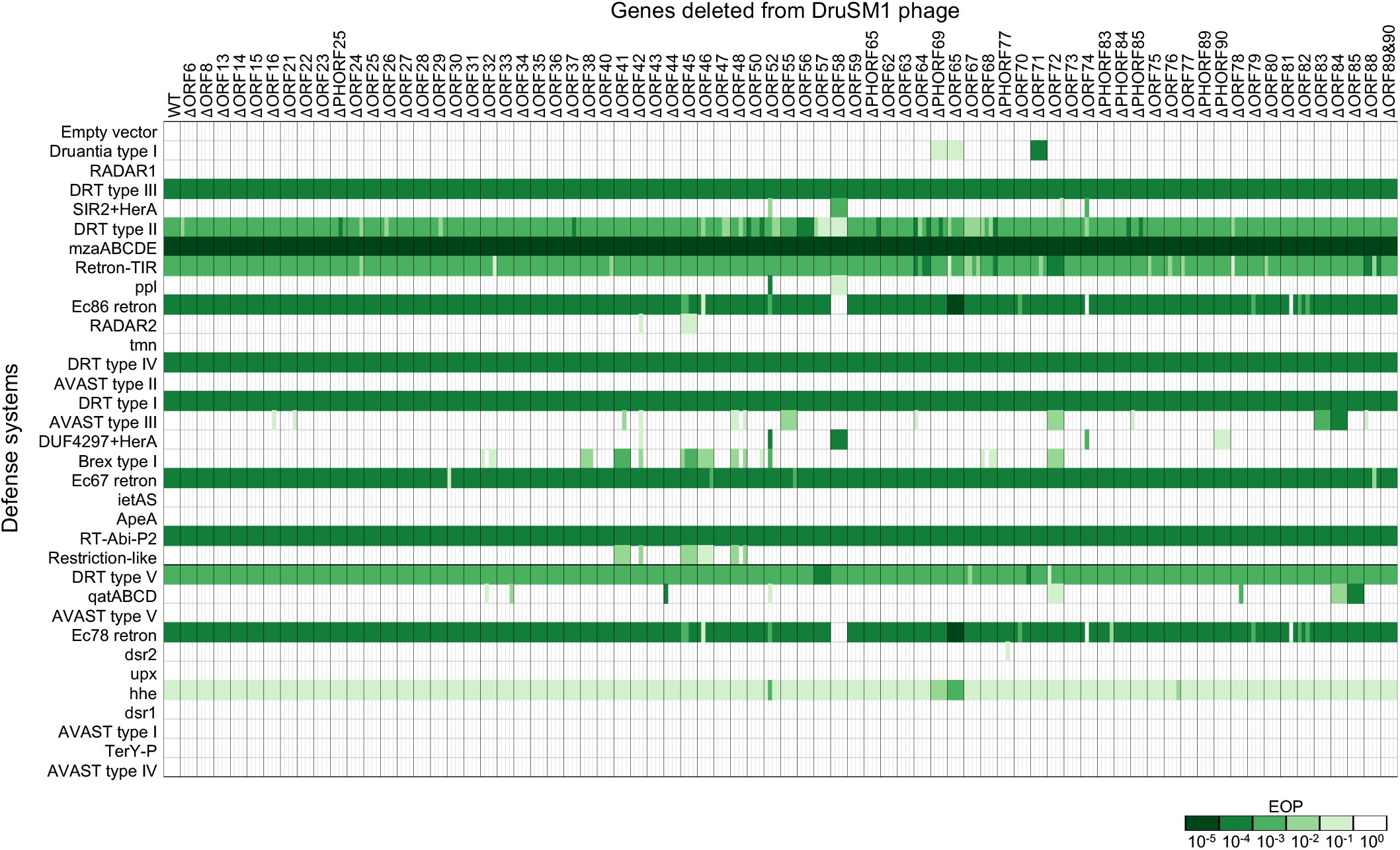
Genetic deletions of phage alter susceptibility to bacterial defense systems (A) Measurement of phage infectivity to bacteria expressing each defense system using the spot assay. Phages with deletions of 73 ORFs infected bacteria harboring defense systems. Four independently synthesized phages were used.

The phages losing the ability to escape from Druantia-defense were ORF71 and ORF65 deletion mutants. The ORF71 deletion mutant showed a significantly decreased EOP compared with that of ORF65 in Druantia type I-bearing strains (*p < 0*.*001*) (Figure 2A and Table 1). Nine ORF-deletion mutants were less infectious to bacteria with Brex1 type I. In particular, gene deletion phages in the “moron, auxiliary metabolism gene and host takeover” region on ORFs 41–48 exhibited reduced infectivity not only against Brex type I but also against restriction-like defense systems (Figure 1C and S1B). Both Brex1 type I and restriction-like are long gene defense systems utilizing ATPase and methylase^7^; thus, an antidefense system targeting these common domains in the ORF41–48 region is expected. Phages with reduced infectivity in bacteria possessing the AVAST type III defense system were deletion mutants of ORF55, ORF72, ORF83, and ORF84. The mutant with the most reduced infectivity was the ORF84 deletion mutant, with a 10^−4^ reduction in EOP. All phages with reduced infectivity against AVAST type III-bearing bacteria formed small plaques. This suggests that the proliferation of the phages was reduced by the AVAST type III defense system. Phages with reduced infectivity against the qatABCD defense system were ORF84 and ORF85 deletion mutants, resulting in reduced EOP by 10^−2^ and 10^−4^, respectively. The phage with reduced infectivity against bacteria harboring the hhe defense system was an ORF65 deletion mutant, exhibiting a decrease in EOP by 10^−3^ and smaller plaque sizes than those of the wild type (Figure S2A). Deletion of PHORF69, located upstream of ORF65, also reduced EOP against hhe-bearing bacteria. Interestingly, ORF58 deletion mutants showed reduced EOP in bacteria expressing SIR2+HerA, DUF4297+HerA, and ppl; however, they showed increased EOP in bacteria harboring Retron-Ec86, Retron-Ec78, and DRT type II, which contain reverse transcriptase domains.

Candidate antidefense systems, deletions of which were expected to result in reduced EOP, were identified by screening a gene deletion library (Figure 2). To confirm that these genes act as antidefense systems, bacteria were prepared through plasmid complementation of the candidate genes, infected with each deletion mutant phage, and the EOP was measured (Figure 3A). Ectopic expression of ORF71 from ΦDruSM1 restored the infectivity of the ORF71 deletion mutant against Drunatia type I, resulting in the formation of same size plaques as those of the wild type (Figure 3A, S2A). Coexpression of Brex1 type I, ORF46, and ORF72 restored the infectivity of each ORF deletion mutant (Figure 3A, S2A). Deletion mutants of ORFs 41, 45, and 48 also showed reduced infectivity in Brex type I-bearing bacteria; however, coexpression of these ORFs with Brex type I did not restore infectivity. The ORF58 deletion mutants showed increased infectivity against bacteria harboring retron-Ec86, retron-Ec78, and DRT type II (Figure 2 and Table 1). These results suggested that ORF58 activates retron-Ec86, retron-Ec78, and DRT type II. Moreover, induced expression of ORF58 caused cytotoxicity in retron-Ec86-bearing strains (Figure 3B), suggesting that ORF58 induces activation of retron-Ec86 and abortive infection. In contrast, ORF58 did not induce cytotoxicity in retron-Ec78-or DRT type II-bearing strains (Fig. S2B), suggesting that factors other than ORF58 are required for retron-Ec78 or DRT type II toxicity.

**Figure 3.**
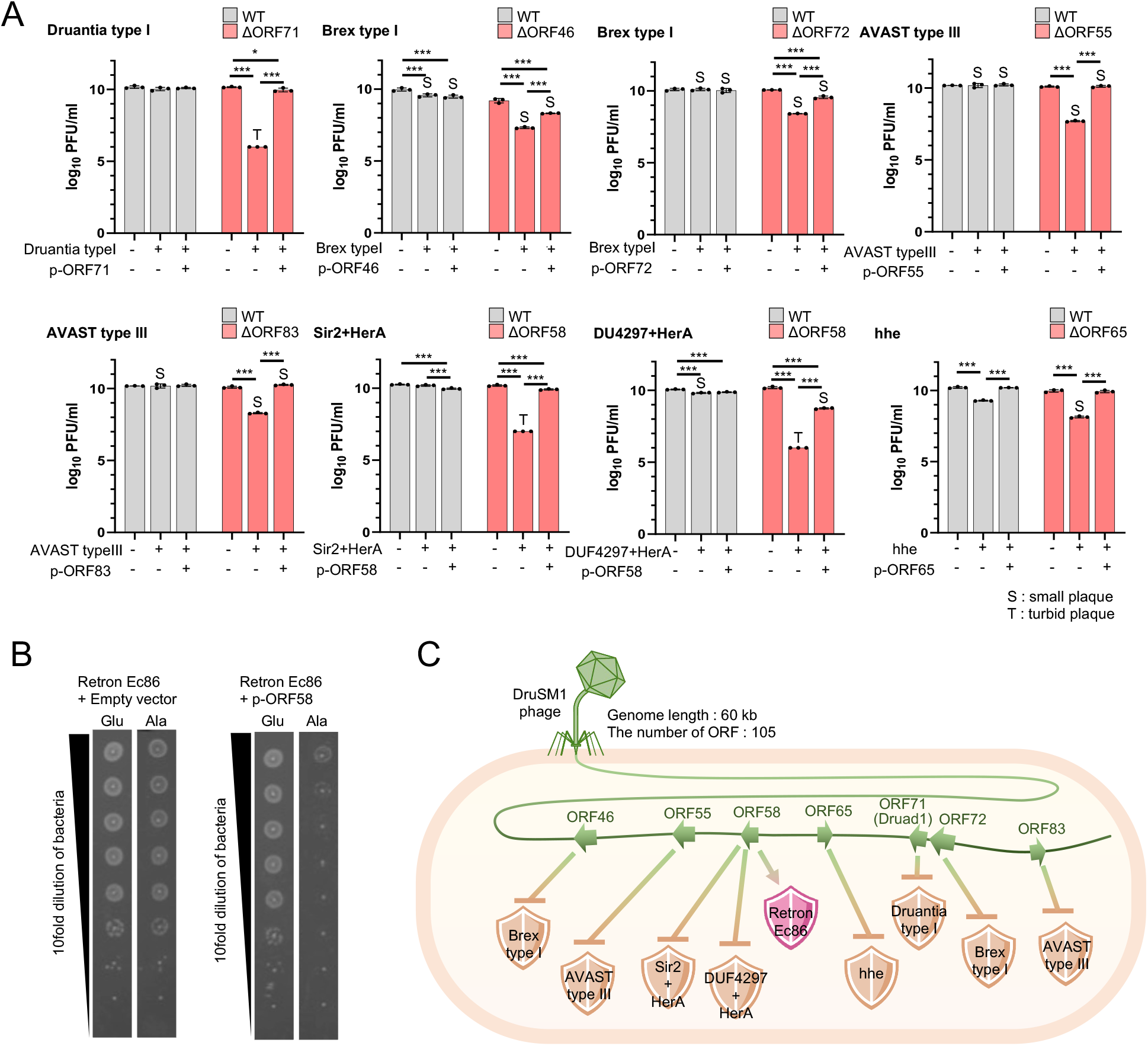
Identification of phage genes modulating susceptibility to defense systems (A) Candidate ORFs identified in the experiment from Figure 2 were cloned and coexpressed with the defense system in *E. coli* DH10B. Phages with deleted ORFs were used for infection followed by plaque counting (N = 3). “T” indicates turbid plaques, whereas “S” indicates reduced plaque size compared with that in the absence of the defense system. (B) *E. coli* DH10B transformed with Retron Ec86-expressing plasmid and arabinose-inducible ORF58-expressing plasmid were cultured on arabinose-containing medium. (C) Schematic diagram depicting the action of the antidefense systems carried by ΦDruSM1 phage. The defense systems highlighted in orange are inhibited by phage genes. The ones highlighted in pink are activated by phage genes.

### Prediction of proteins that inactivate and activate defense systems

Functional prediction of antidefense or activator genes was conducted by performing a domain search using the HHpred server, with PFAM and COG_KOG serving as the target databases (Table 2 and 3). ORF46, exhibiting anti-Brex type I activity, was predicted to be a “Trimethylamine methyltransferase corrinoid protein”. ORF55, an ORF with anti-AVAST type III activity, was predicted to be a “ATP-dependent DNA ligase”. ORF58, which has anti-SIR2+HerA and DUF4297+HerA activity as well as Reron Ec86 sensor activity was predicted to be a “Mu-like prophage host-nuclease inhibitor protein Gam”. ORF65, showing anti-hhe activity was predicted to be a “Transcriptional regulator protein (SplA) “. ORF71, exhibiting anti-Druantia type I activity, was predicted to belong to a “Family of unknown function (DUF6614) “. ORF72, an ORF with anti-Brex type I activity was predicted to be a “KfrA_N; Plasmid replication region DNA-binding N-term”. Finally, ORF83, exhibiting anti-AVAST type III activity was predicted to be a “Smf; Predicted Rossmann fold nucleotide-binding protein DprA/Smf involved in DNA uptake”.

**Table 2.** Anti-defense genes discovered from DruSM1 phage.

|  |  |  |  |  |  |
| --- | --- | --- | --- | --- | --- |
| ORF46 | Brex type I | 62 AA | Trimethylamine methyltransferase corrinoid protein | 76.55 | 9.4 |
| ORF55 | AVAST type III | 304 AA | ATP-dependent DNA ligase | 100 | 2.7E-38 |
| ORF58 | Sir2+HerA, DUF4297+HerA | 152 AA | Mu-like prophage host-nuclease inhibitor protein Gam | 97.02 | 0.059 |
| ORF65 | Hhe | 74 AA | Transcriptional regulator protein (SplA) | 72.24 | 8.1 |
| ORF71 (Druad1) | Druantia type I | 44 AA | Family of unknown function (DUF6614) | 40.55 | 17 |
| ORF72 | Brex type I | 204 AA | KfrA_N; Plasmid replication region DNA-binding N-term | 64.96 | 39 |
| ORF83 | AVAST type III | 130 AA | Smf; Predicted Rossmann fold nucleotide-binding protein DprA/Smf involved in DNA uptake | 99.81 | 5E-18 |

**Table 3.** Defense activator genes discovered from DruSM1 phage.

|  |  |  |  |  |  |
| --- | --- | --- | --- | --- | --- |
| ORF58 | Retron Ec86 | 152 AA | Mu-like prophage host-nuclease inhibitor protein Gam | 97.02 | 0.059 |

### Phages equipped with Druad1 can infect bacteria possessing Druantia type I

In this study, we focused on Druantia type I because this defense system exhibited the most extensive defense activity in our collection of 263 *E. coli* phage libraries (Figure 4A and B). Among the antidefense genes identified in this study, we selected ORF71, which inhibits Druantia type I. ORF71 was named Druad1 (Druantia antidefense 1) because of its ability to inhibit Druantia type I. Other members of the Quenovirinae family to which ΦDruSM1 belonged included KSA3, KSA8, KSS4, KSW4, and SHIN8 (Figure 4C)^21,22^. Of these six phages, only ΦDruSM1 killed DH10B expressing Druantia type I (Figure 4D and Figure S3A). Notably, ΦDruSM1ΔDruad1 failed to block Druantia type I, indicating the indispensable role of Druad1 in the inactivation of Druantia type I during phage infection.

**Figure 4.**
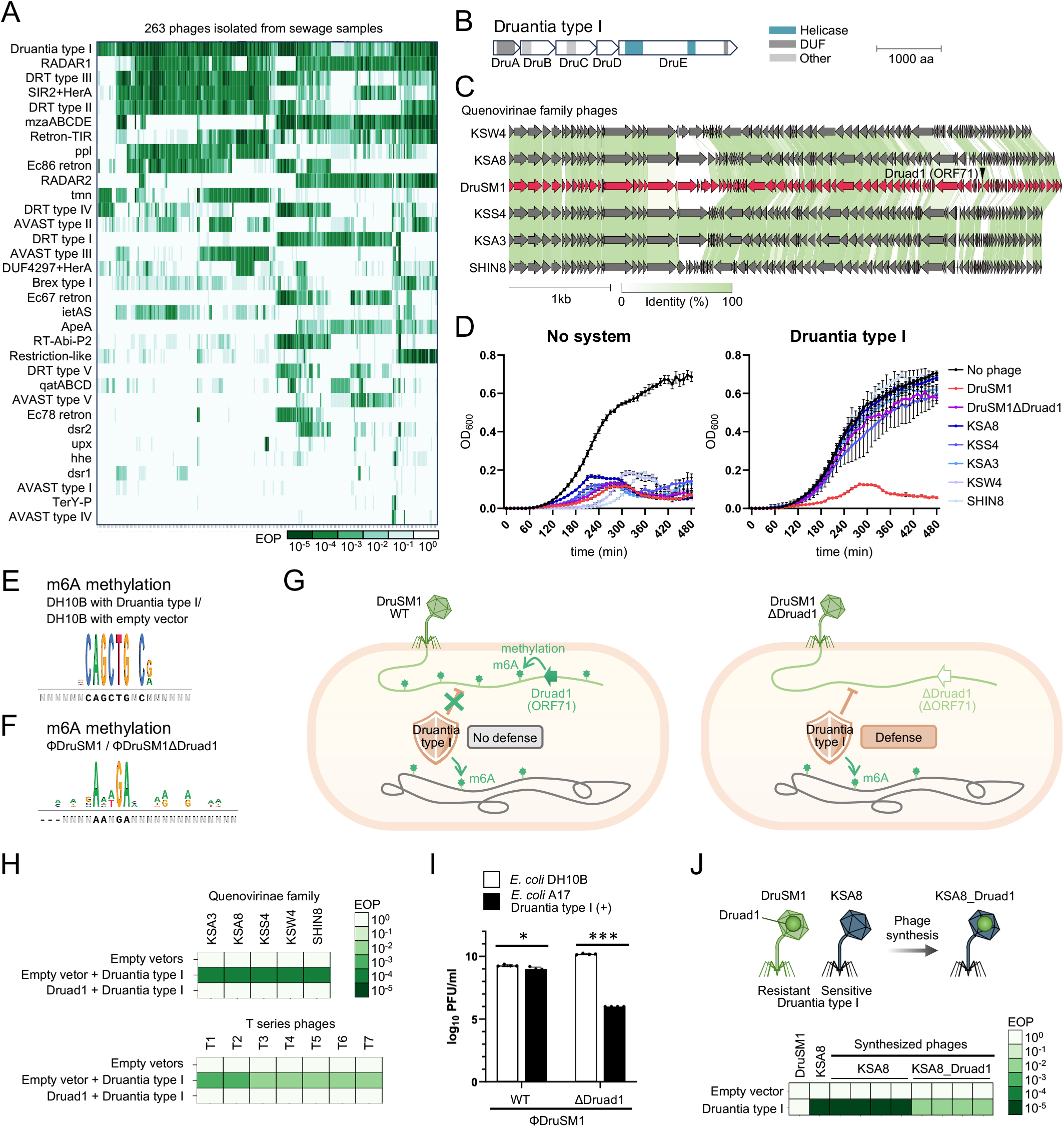
Druad1 suppresses the potent defense system Druantia type I (A) 263 sewage-derived *E. coli* phages infected *E. coli* DH10B possessing 33 different antiphage defense systems, and plaque formation efficiency was calculated. The defense systems are listed in order of decreasing plaque formation efficiency. (B) Gene structure of Druantia type I^7^. (C) Genome comparison of phages belonging to the Quenovirinae seuratvirus family isolated in this study. (D) Comparison of bactericidal activity of Quenovirinae phages and Druad1 (ORF71) deletion mutant of ΦDruSM1. Phages infected DH10B expressing Druantia type I at MOI 0.01, and bacterial growth was measured. (E) Unique m6A-modified DNA sequence found in DH10B with Druantia Type I and absent in DH10B with empty vector. (F) Unique m6A-modified DNA sequence found in ΦDruSM1 and absent in ΦDruSM1ΔDruad1. (G) A schematic diagram illustrating the mechanism by which Druad1-bearing phages evade the Druantia type I defense system. (H) Defense activity of Druantia type I in DH10B expressing Druad1. (I) Phage sensitivity of *E. coli* DH10B and A17 were measured using the spot assay. (J) Druad1 was artificially inserted into the ΦKSA8 genome, resulting in ΦKSA8_Druad1. Synthesized phages infected *E. coli* DH10B harboring Druantia type I antiphage defense systems. Phage infectivity was measured using the spot assay.

We then searched for homologs of Druad1 and found no proteins with a BLAST value <0.1, suggesting that it is an extremely rare gene. Furthermore, the genome of ΦDruSM1 showed 83.0 % homology with that of the most closely related phage, vB_Ecos_SA126VB, indicating that ΦDruSM1 itself is a novel phage (Figure S4A and B). Druantia type I system harbors a DNA helicase domain (Figure 4B). Given that type I restriction-modification systems, which encode helicase domain proteins, facilitate DNA translocation upon recognizing unmethylated restriction sites, we speculated that Druantia type I employs DNA methylation as well.^23^ Utilizing PacBio sequencing, the methylation status of DNA in strains expressing Druantia type I was compared with that in non-expressing strains. As a result, methylation of m6A in the CAGCTGNC sequence was only observed in strains expressing Druantia type I (Figure 4E), suggesting that Druantia type I adds m6A methylation to the host bacterial genome. Considering the potential involvement of DNA methylation in the function of Druad1, the methylation status of DNA in ΦDruSM1 and ΦDruSM1ΔDruad1 was compared. Consequently, m6A methylation in the AANGA sequence was confirmed only in ΦDruSM1 carrying Druad1 (Figure 4F). Thus, while Druantia type I distinguishes between bacterial genomic DNA and phage DNA through m6A methylation, Druad1 may facilitate evasion from recognition by Druantia type I by inducing m6A modification in phage DNA (Figure 4G). Various phages, including Quenovirinae family and T-series, infected *E. coli* expressing both Druantia type I and Druad1, but not *E. coli* expressing Druantia type I alone (Figure 4H). This suggests that in the presence of sufficient Druad1 expression, phages are capable of escaping detection by Druantia type I. To confirm the activity of Druad1 against native Druantia type I, the infectivity of ΦDruSM1ΔDruad1 was tested using the *E. coli* clinical isolate A17 harboring Druantia type I. Sequence alignment of the Druantia type I gene from Gao et al. and the Druantia type I gene from *E. coli* A17 showed an overall similarity of more than 98 %. The similarities for each gene were 99.2 %, 98.5 %, 98.4 %, 99.7 %, and 99.4 % for DruA, DruB, DruD, and DruE, respectively (Figure S3B). ΦDruSM1 infected *E. coli* A17 harboring Druantia; however, the infectivity of ΦDruSM1ΔDruad1 was markedly reduced, suggesting that Druad1 also functions against clinical isolates harboring Druantia type I (Fig. 4I). In phage therapy, phages that can evade the potent Druantia type I defense system are valuable. Therefore, we decided to artificially create a phage that can evade the Druantia type I defense system: ΦKSA8, which is similar to ΦDruSM1 and lacks the Druad1 homolog. Incorporating Druad1 into this phage increased its infectivity against DH10B expressing Druantia type I (Figures 4J and S3C).

## Discussion

In this study, we created a phage knockout library using ΦDruSM1 with a genome size of 60 kb. A comprehensive assay, combining the phage knockout library with the defense system expression vector library, revealed that ΦDruSM1 possesses more than seven antidefense systems.

For creating knockout strains of ΦDruSM1, we used in vitro phage synthesis by assembling PCR fragments (Figure 1A)^17,18^. Although the genome length for phage synthesis in vitro was reported in 2023 to be approximately 50 kb^18^, we succeeded in artificially synthesizing a 60 kb phage, ΦDruSM1, and knocked out 73 out of 105 genes (Figure 1B). In the reboot experiments using gene knockout phages, deleted genes resulting in the formation of 10 or fewer plaques, were classified as essential, whereas those leading to the formation of 11 or more plaques were classified as nonessential; however, this is not a perfect classification. This is because genes essential for phage proliferation within HST08 strain used in this study are not always essential for proliferation within other *E coli* strains. For instance, if DruSM1 harbors an inhibitor against the defense system of HST08, this inhibitor may be essential for proliferation within HST08 but not necessarily essential within other *E. coli* strains. This study implemented a knockout library using the ΦDruSM1 phage infecting *E. coli*. As phage in vitro synthesis methods and genetic engineering methods have advanced, this approach will be applicable to other phages infecting diverse bacterial species in the future. Large-genome phages such as jumbo phages are believed to have special antidefense systems^24^; however, a synthesis method for jumbo phages has not yet been established.

Although many antiphage defense systems have been discovered, systems that counteract them have been rarely reported^3^. On average, a bacterium has been reported to have at least five defense systems^2,25-28^, and phages have likely evolved the means to counteract them. In this study, we identified more than seven antidefense systems in a single phage (Figure 3A). Considering that more than 100 defense systems have already been reported^2^ and many more subspecies exist, we assumed that ΦDruSM1 has a greater number of antidefense systems. As many anti-defense systems were found in ΦDruSM1 alone, it is expected that many more antidefense systems will be discovered in the future. Many of the genes identified as antidefense systems are hypothetical proteins, which opens up the possibility of identifying the functions of previously unknown phage genes. In some cases, plasmid complementation of the knockout gene did not restore the phenotype of the knockout phage (Figure 2 and 3A). This could be due to a polar effect, in which the knockout gene affects the expression of surrounding genes. This could also be due to inadequate phage annotation because phages often encode small proteins^29^. In addition, as phage-derived RNAs are also known to inhibit these defense systems^1,4^, it is possible that noncoding nucleic acids rather than proteins were responsible for these results.

Notably, our experiments also revealed that the Gam protein of ΦDruSM1 acts as an activator of Retron Ec86 (Figure 3B). This finding aligned with previous studies on Gam in λ phage serving as a sensor for Retron^27^, further supporting the validity of our methodology. Of note, Gam simultaneously inhibited Sir2 + HerA and DUF4297 + HerA, suggesting the inhibition of the common helicase domain known as HerA; however, the mechanism remains unclear (Tables 2 and 3). One gene may be a sensor for another, such as Ocr, which acts as an anti-RM or anti-Brex system and is sensed by PARIS^2,30^, and Gam, like Ocr, may be involved in various antidefense systems. More than seven genes were found to inhibit the bacterial defense systems; however, only one gene was identified as activator of the defense systems. One reason for this may be that many defense system activator genes are essential genes^2,27,31^. Because essential genes in phages cannot be genetically deleted, searching for activator genes of the defense system using our screening method is difficult.

It has been revealed that both Druantia type I and Druad1 are involved in DNA methylation at m6A sites (Figure 4E-G). However, within Druantia type I, which lacks a Methylase domain, the gene responsible for m6A methylation has yet to be identified.^20,32^ Similarly, Druad1, a small 44-amino acid gene, also lacks a Methylase domain, leaving the mechanism of m6A site methylation unclear. Considering that DNA adenine methyltransferase and DNA cytosine methyltransferase in *E. coli* MG1655 consist of 278 and 472 amino acids,^33^ respectively, it is unlikely that Druad1 acts alone in methylation. Instead, its interaction with other factors suggests a potential collaborative role in methylation. Furthermore, since Druad1 neutralizes the defense of Druantia type I against phages beyond its original host ΦDruSM1 (Figure 4H), Druad1 may be associated with host methyltransferases.

The in vitro synthesis system of phages that we utilized in this study has limited efficiency in synthesizing phages with large genomes^34,35^. Therefore, employing the same methodology to identify antidefense genes from phages with large genomes presents challenges. However, applying methods such as random mutagenesis to large-genome phages may enable the construction of a more diverse library of gene knockouts, potentially leading to the discovery of a greater number of antidefense genes. Additionally, in this study, we explored antidefense genes by expressing defense systems in laboratory strains of *E. coli* using plasmids. However, overexpression of genes by plasmids may not fully reflect native conditions and physiological conditions of bacteria. Conducting large-scale studies infecting phage knockout libraries to various clinical isolates in the future would elucidate the interactions and evolution of diverse defense genes and antidefense genes under native conditions.

Owing to the escalating problem of drug-resistant bacteria, phage-based antibacterial therapy is gaining attention^18,36-40^. As phage infectivity is defined by the defense system and bacterial receptor affinity, phage therapy requires screening for phages that can efficiently kill clinical isolates from the environment or library^18,21,39,41^. Incorporation of antidefense genes into phages can facilitate the implementation of phage therapy without the limitations imposed by the defense system. Our study demonstrated that phages artificially incorporating Druad1 killed bacteria harboring Druantia type I (Figure 4J and S3C). We believe that this set of methods provides a roadmap for enhancing the host range and bactericidal effects of phages during phage therapy.

## Acknowledgments

We thank Dr. Nishimasu Hiroshi and Mr. Junichiro Ishikawa from Tokyo University for the fruitful discussion during our manuscript preparation. Funding: This work was supported by the Japan Agency for Medical Research and Development (Grant No. 23wm0325065, 22fk0108532, 24fk0108698 and JP21gm1610002 to K. Kiga), and JSPS KAKENHI (Grant Nos. 21H02110 to K. Kiga, and 22K20575 to SO). The funders had no role in the study design, data collection and analysis, decision to publish, or preparation of the manuscript.

## Author contributions

SO designed and conducted the experiments, analyzed the data, and drafted the manuscript. AHA conducted experiments and drafted the manuscript. K. Kondo, WN, SW, KC, TN and BZ provided expertise in bioinformatics analysis. AT, YW, and YS conducted the experiments and contributed to data collection. MS, YT, and KW critically reviewed the manuscript. K. Kiga designed and supervised the study, provided funding, and drafted and approved the final version of the manuscript.

## Supplementary Figure Legends

**Figure S1**. Confirmation of each DruSM1 open reading frame (ORF) deletion mutant, and coding sequence map containing the antidefense and defense sensor genes of ΦDruSM1, relating to Figure 1.

(A) Photographs of plaque PCR electrophoresis for each ORF deleted mutant resulting in altered defense activity. (B) Coding sequence map of ΦDruSM1 containing antidefense activity and defense sensor genes on synthetic efficiency of ORF deletion mutants. Descriptions of defense with altered activity in the Figure 1C map were added here.

**Figure S2**. Photographic data on antidefense or defense sensor activity, relating to Figure 3.

(A) Photographic data of spot assays evaluating antidefense activity. ΦDruSM1 WT and respective ΦDruSM1 ORF deletion mutant were spotted onto *E. coli* DH10B harboring respective defense systems and antidefense genes. (B) Photographic data of toxicity assay on combinations for retron Ec78 and ORF58, or DRT type 2 and ORF58. *E. coli* DH10B harboring respective defense systems and ORF58 were grown in LB supplemented with glucose, and a 10-fold dilution of each O/N culture was made and spotted onto glucose- or arabinose-supplemented LB plates.

**Figure S3**. Support data for Druantia type I analysis, relating to Figure 4.

(A) Defense pattern of 6 Quenovirinae phages in the defense library by Gao et al. ΦDruSM1, ΦSHIN8, ΦKSS4, ΦKSA3, ΦKSW4, and ΦKSA8 were screened using the spot assay in *E. coli* DH10B harboring pLG001-034. The fold reduction in EOP was calculated based on the EOP on DH10B harboring pLG001 (no defense system). (B) Comparison of Drunatia type I between the defense library by Gao et al. and *E. coli* clinical isolate A17 strain. Coding sequence (CDS) annotations were done using PADLOC and CDS alignment was done using Clinker. (C) Photographic data of the bactericidal effect of ΦKSA8 artificially expressing the anti-Druantia type I gene following infection of bacteria expressing Druantia type I using the spot assay.

**Figure S4**. Phage classification of ΦDruSM1, relating to Figure 4.

(A) Phylogenetic tree of ΦDruSM1 and similar phages. Viptree was used for constructing the protein-based phylogenic tree. (B) Phages with similar nucleotide identity with ΦDruSM1. The genomes of phages showing similar nucleotide identity were identified using online blast, while average nucleotide identity (ANI) was determined using VIRIDIC with default settings.

## Notes

### Competing Interest Statement

SO, AHA, YW, YT, KW, and K. Kiga. are coinventors of a patent submitted by the National Institute of Infectious Diseases based on the results reported in this paper.

