## Supplementary material for "Systematic Discovery of Phage Genes that Inactivate Bacterial Immune Systems": Ojima et al. Supplementary Figures.pdf

### Figure S1

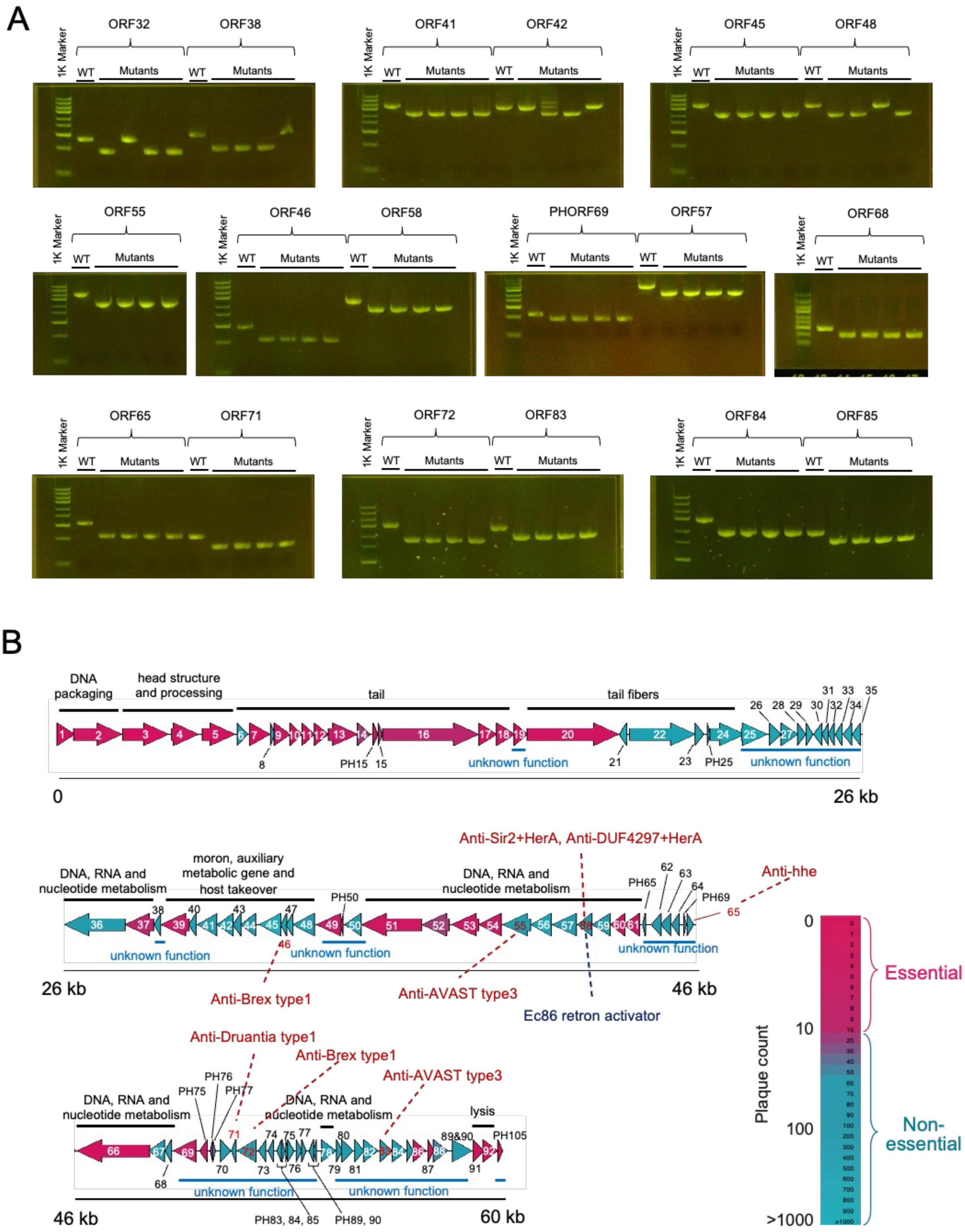
A
1K Marker
ORF32
ORF38
ORF41
ORF42
ORF45
ORF48
ORF55
ORF46
ORF58
PHORF69
ORF57
ORF68
ORF65
ORF71
ORF72
ORF83
ORF84
ORF85
WT
Mutants
B
DNA packaging
head structure and processing
tail
tail fibers
unknown function
0
26 kb
DNA, RNA and nucleotide metabolism
moron, auxiliary metabolic gene and host takeover
PH50
PH65
PH69
Anti-Sir2+HerA, Anti-DUF4297+HerA
Anti-hhe
Anti-Brex type1
Anti-AVAST type3
Ec86 retron activator
46 kb
Anti-Druantia type1
PH75
PH76
PH77
lysis
PH105
PH83, 84, 85
PH89, 90
60 kb
Plaque count
0
10
100
>1000
Essential
Non-essential

### Figure S2

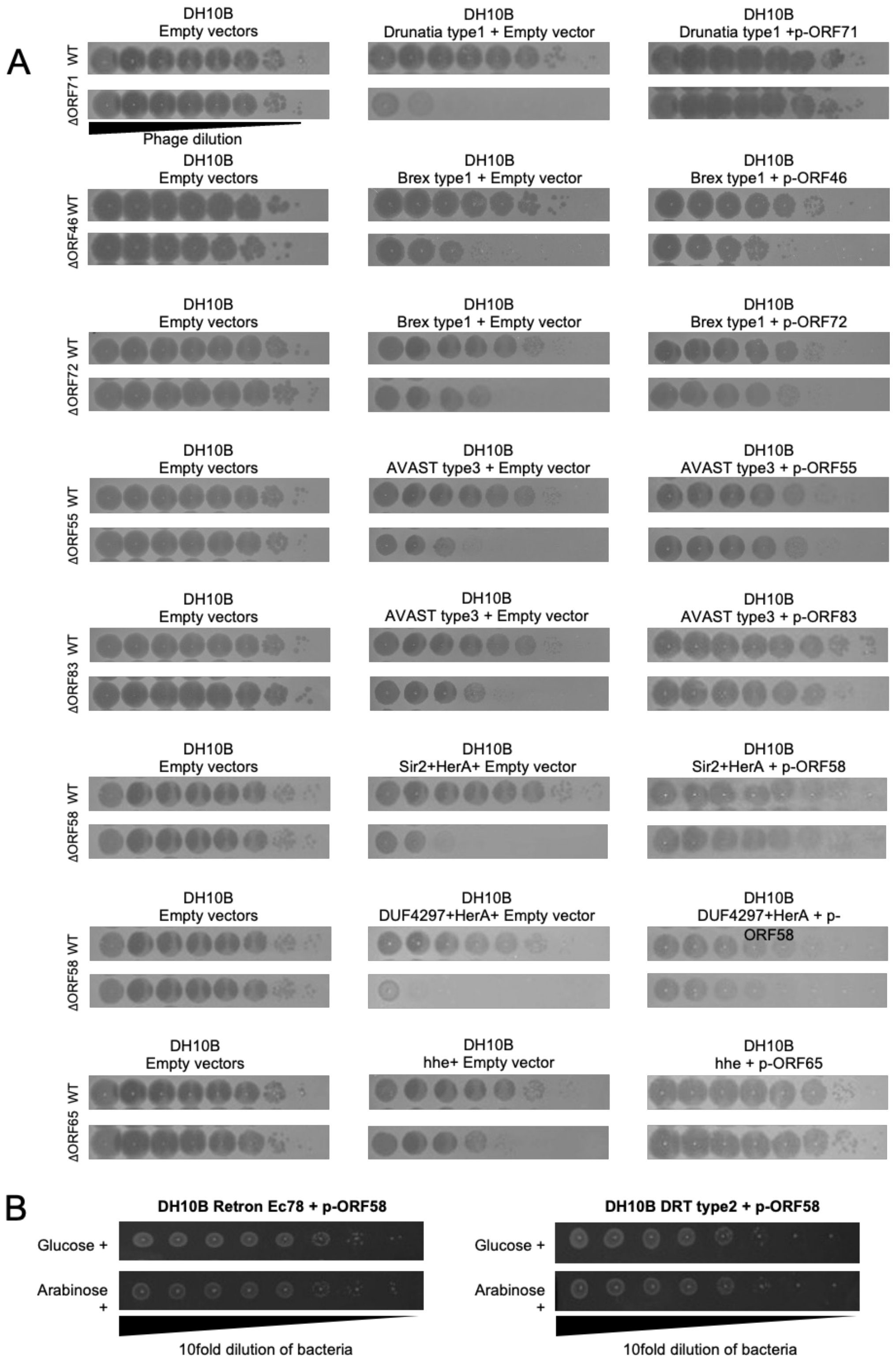
A
DH10B Empty vectors
DH10B Drunatia type1 + Empty vector
DH10B Drunatia type1 +p-ORF71
WT
ΔORF71
Phage dilution
DH10B Empty vectors
DH10B Brex type1 + Empty vector
DH10B Brex type1 + p-ORF46
WT
ΔORF46
DH10B Empty vectors
DH10B Brex type1 + Empty vector
DH10B Brex type1 + p-ORF72
WT
ΔORF72
DH10B Empty vectors
DH10B AVAST type3 + Empty vector
DH10B AVAST type3 + p-ORF55
WT
ΔORF55
DH10B Empty vectors
DH10B AVAST type3 + Empty vector
DH10B AVAST type3 + p-ORF83
WT
ΔORF83
DH10B Empty vectors
DH10B Sir2+HerA+ Empty vector
DH10B Sir2+HerA + p-ORF58
WT
ΔORF58
DH10B Empty vectors
DH10B DUF4297+HerA+ Empty vector
DH10B DUF4297+HerA + p-ORF58
WT
ΔORF58
DH10B Empty vectors
DH10B hhe+ Empty vector
DH10B hhe + p-ORF65
WT
ΔORF65
B
DH10B Retron Ec78 + p-ORF58
Glucose +
Arabinose +
10fold dilution of bacteria
DH10B DRT type2 + p-ORF58
Glucose +
Arabinose +
10fold dilution of bacteria

### Figure S3

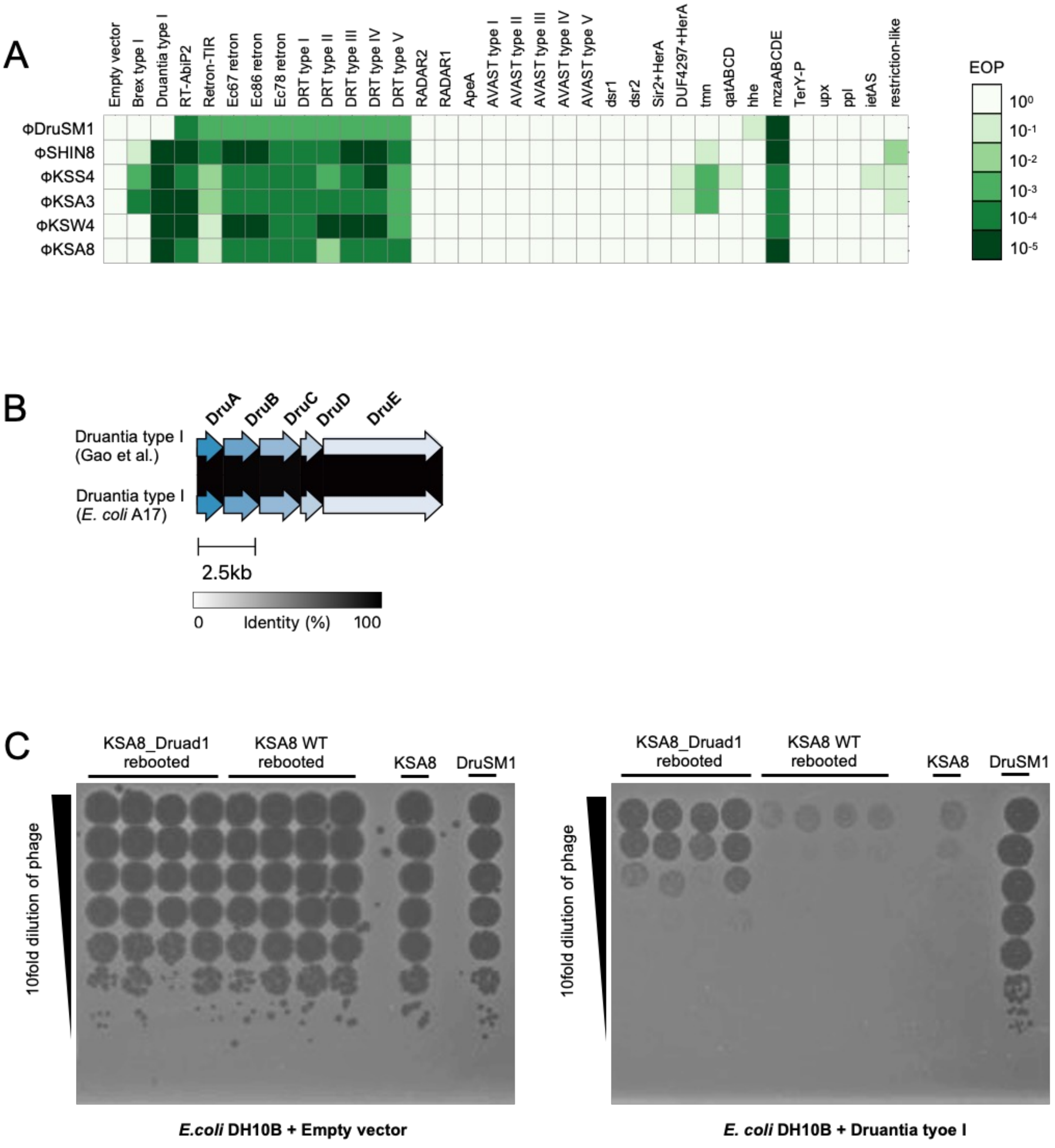
A
Empty vector
Brex type I
Druantia type I
RT-AbiP2
Retron-TIR
Ec67 retron
Ec86 retron
Ec78 retron
DRT type I
DRT type II
DRT type III
DRT type IV
DRT type V
RADAR2
RADAR1
ApeA
AVAST type I
AVAST type II
AVAST type III
AVAST type IV
AVAST type V
dsr1
dsr2
Sir2+HerA
DUF4297+HerA
tmn
qatABCD
hhe
mzaABCDE
TerY-P
upx
ppl
ietAS
restriction-like
ΦDruSM1
ΦSHIN8
ΦKSS4
ΦKSA3
ΦKSW4
ΦKSA8
EOP
10⁰
10⁻¹
10⁻²
10⁻³
10⁻⁴
10⁻⁵
B
DruA
DruB
DruC
DruD
DruE
Druantia type I (Gao et al.)
Druantia type I (E. coli A17)
2.5kb
0
Identity (%)
100
C
KSA8_Druad1 rebooted
KSA8 WT rebooted
KSA8
DruSM1
10fold dilution of phage
E.coli DH10B + Empty vector
E. coli DH10B + Druantia tyoe I

Figure S4

A

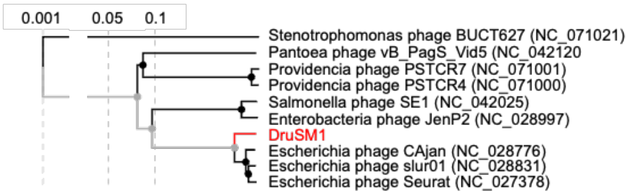
0.001
0.05
0.1
Stenotrophomonas phage BUCT627 (NC_071021)
Pantoea phage vB_PagS_Vid5 (NC_042120
Providencia phage PSTCR7 (NC_071001)
Providencia phage PSTCR4 (NC_071000)
Salmonella phage SE1 (NC_042025)
Enterobacteria phage JenP2 (NC_028997)
DruSM1
Escherichia phage CAjan (NC_028776)
Escherichia phage slur01 (NC_028831)
Escherichia phage Seurat (NC_027378)

B

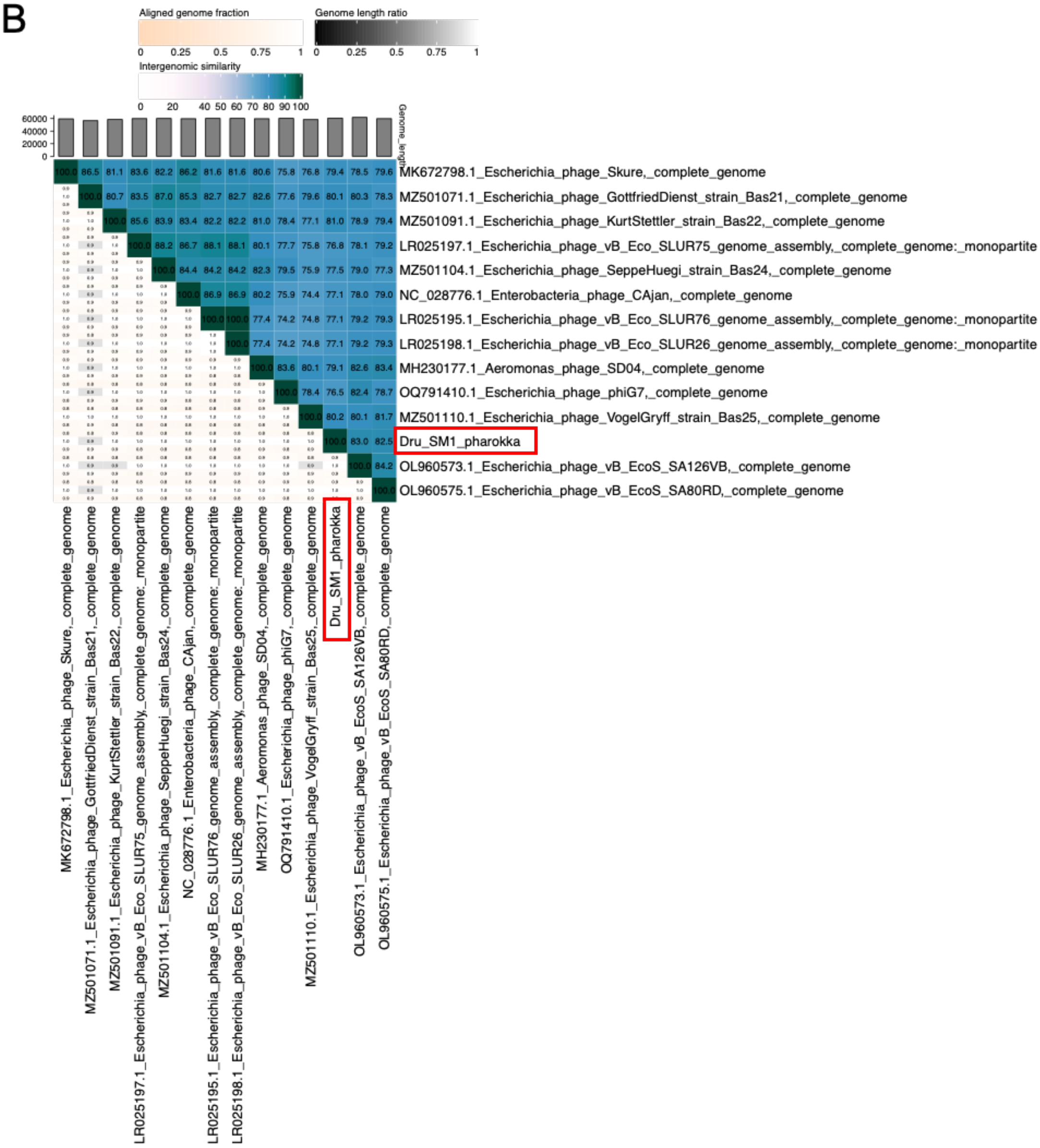
Aligned genome fraction
0
0.25
0.5
0.75
1
Genome length ratio
0
0.25
0.5
0.75
1
Intergenomic similarity
0
20
40 50 60 70 80 90 100
60000
40000
20000
0
Genome_length
MK672798.1_Escherichia_phage_Skure,_complete_genome
MZ501071.1_Escherichia_phage_GottfriedDienst_strain_Bas21,_complete_genome
MZ501091.1_Escherichia_phage_KurtStettler_strain_Bas22,_complete_genome
LR025197.1_Escherichia_phage_vB_Eco_SLUR75_genome_assembly,_complete_genome:_monopartite
MZ501104.1_Escherichia_phage_SeppeHuegi_strain_Bas24,_complete_genome
NC_028776.1_Enterobacteria_phage_CAjan,_complete_genome
LR025195.1_Escherichia_phage_vB_Eco_SLUR76_genome_assembly,_complete_genome:_monopartite
LR025198.1_Escherichia_phage_vB_Eco_SLUR26_genome_assembly,_complete_genome:_monopartite
MH230177.1_Aeromonas_phage_SD04,_complete_genome
OQ791410.1_Escherichia_phage_phiG7,_complete_genome
MZ501110.1_Escherichia_phage_VogelGryff_strain_Bas25,_complete_genome
Dru_SM1_pharokka
OL960573.1_Escherichia_phage_vB_EcoS_SA126VB,_complete_genome
OL960575.1_Escherichia_phage_vB_EcoS_SA80RD,_complete_genome

#### Supplementary Figure Legends

**Figure S1.** Confirmation of each DruSM1 open reading frame (ORF) deletion mutant, and coding sequence map containing the antidefense and defense sensor genes of ΦDruSM1, relating to Figure 1.
(A) Photographs of plaque PCR electrophoresis for each ORF deleted mutant resulting in altered defense activity. (B) Coding sequence map of ΦDruSM1 containing antidefense activity and defense sensor genes on synthetic efficiency of ORF deletion mutants. Descriptions of defense with altered activity in the Figure 1C map were added here.

**Figure S2.** Photographic data on antidefense or defense sensor activity, relating to Figure 3.
(A) Photographic data of spot assays evaluating antidefense activity. ΦDruSM1 WT and respective ΦDruSM1 ORF deletion mutant were spotted onto *E. coli* DH10B harboring respective defense systems and antidefense genes. (B) Photographic data of toxicity assay on combinations for retron Ec78 and ORF58, or DRT type 2 and ORF58. *E. coli* DH10B harboring respective defense systems and ORF58 were grown in LB supplemented with glucose, and a 10-fold dilution of each O/N culture was made and spotted onto glucose- or arabinose-supplemented LB plates.

**Figure S3.** Support data for Druantia type I analysis, relating to Figure 4.
(A) Defense pattern of 6 Quenovirinae phages in the defense library by Gao et al. ΦDruSM1, ΦSHIN8, ΦKSS4, ΦKSA3, ΦKSW4, and ΦKSA8 were screened using the spot assay in *E. coli* DH10B harboring pLG001-034. The fold reduction in EOP was calculated based on the EOP on DH10B harboring pLG001 (no defense system). (B) Comparison of Drunatia type I between the defense library by Gao et al. and *E. coli* clinical isolate A17 strain. Coding sequence (CDS) annotations were done using PADLOC and CDS alignment was done using Clinker. (C) Photographic data of the bactericidal effect of ΦKSA8 artificially expressing the anti-Druantia type I gene following infection of bacteria expressing Druantia type I using the spot assay.

**Figure S4.** Phage classification of ΦDruSM1, relating to Figure 4.
(A) Phylogenetic tree of ΦDruSM1 and similar phages. Viptree was used for constructing the protein-based phylogenic tree. (B) Phages with similar nucleotide identity with ΦDruSM1. The genomes of phages showing similar nucleotide identity were identified using online blast, while average nucleotide identity (ANI) was determined using VIRIDIC with default settings.
